# Deep brain stimulation modulates directional frontolimbic connectivity in obsessive‐compulsive disorder

**DOI:** 10.1101/809269

**Authors:** Egill Axfjord Fridgeirsson, Martijn Figee, Judy Luigjes, Pepijn van den Munckhof, P.Richard Schuurman, Guido van Wingen, Damiaan Denys

## Abstract

Deep brain stimulation (DBS) is effective for patients with treatment‐refractory obsessive‐compulsive disorder. DBS of the ventral anterior limb of the internal capsule (vALIC) rapidly improves mood and anxiety with optimal stimulation parameters. To understand these rapid effects of vALIC‐DBS, we studied functional interactions within the affective amygdala circuit. We compared resting state functional magnetic resonance imaging data during chronic stimulation versus one week of stimulation discontinuation in patients, and obtained two resting state scans from matched healthy volunteers to account for test‐retest effects. Imaging data were analyzed using functional connectivity analysis and dynamic causal modelling. Improvement in mood and anxiety following DBS was associated with reduced amygdala‐insula functional connectivity. Directional connectivity analysis revealed that DBS increased the impact of the ventromedial prefrontal cortex on the amygdala, and decreased the impact of the amygdala on the insula. These results highlight the importance of the amygdala circuit in the pathophysiology of OCD, and suggest a neural systems model through which negative mood and anxiety are modulated by vALIC‐DBS for OCD and possibly other psychiatric disorders.

**One Sentence Summary:** Deep brain stimulation improves mood and anxiety in obsessive‐compulsive disorder by altering connectivity between the amygdala, insula and prefrontal cortex.

## Introduction

Obsessive‐compulsive disorder (OCD) is a psychiatric disorder with an estimated lifetime prevalence of 2% in the general population ^1,2^. The main symptoms are anxiety, obsessive thoughts (obsessions) and repetitive behaviors (compulsions). Patients are commonly treated with cognitive behavioral therapy and/or selective serotonin reuptake inhibitors ^3^. Treatment for patients who do not respond sufficiently includes clomipramine and a combination of SSRIs with antipsychotics. Approximately 10% of OCD patients remain treatment refractory and continue to experience symptoms despite pharmacological and behavioral treatment ^3^. For those patients deep brain stimulation (DBS) is an emerging treatment option with approximately 60% responder rate ^4^.

In DBS, electrodes are implanted in brain regions which can then be selectively and focally stimulated with electrical impulses. DBS has been tested as a viable treatment option in several psychiatric conditions such as depression, anorexia nervosa and addiction ^5,6^. In OCD, the most common target regions include striatal regions such as nucleus accumbens (NAc), ventral capsule/ventral striatum and ventral anterior limb of the internal capsule (vALIC). Other common regions are subthalamic nucleus, inferior thalamic peduncle and more recently, medial forebrain bundle ^7,8^. Once stimulation parameters have been optimized, DBS of the vALIC results in a typical sequence of symptom improvements. Patients initially experience rapid improvements of mood and anxiety, followed by more gradual decrease of obsessions and compulsion which may take several weeks and often require additional behavioral therapy for several months ^9,10^. We found that decreased obsessions and compulsions following DBS were associated with normalization of frontostriatal network function ^11^. However, it remains puzzling how vALIC DBS induces its rapid changes in mood and anxiety.

Here we investigated whether rapid mood and anxiety effects of vALIC‐DBS are due to modulation of circuits involving a predominant role of the amygdala. The amygdalae are crucial for the detection of salient events and the initiation of anxiety ^12^. Mood and anxiety disorders have been consistently associated with increased activity of amygdala and insula, and decreased activity of the prefrontal cortex ^13–17^. Functional connectivity between amygdala and insula is positively correlated to anxiety ^18^, whereas functional connectivity between amygdala and ventromedial prefrontal cortex (vmPFC) is negatively correlated to anxiety and negative affect ^19,20^. The vALIC DBS target region is strongly connected with the amygdala, insula, and vmPFC ^21^. We therefore hypothesized that rapid mood and anxiety effects of vALIC DBS result from modulation in connectivity between the amygdala, insula and vmPFC.

In this study, we used two methods to assess changes in connectivity as measured with resting state functional magnetic resonance imaging (fMRI). First, we used functional connectivity to measure correlations in spontaneous slow fluctuations (<0.1Hz) in blood oxygen level dependent (BOLD) signals between the amygdala and the rest of the brain. This method has shown considerable intra‐subject reproducibility ^22,23^ and has been linked to behavioral variability ^24^. Because the amygdala is composed of distinct nuclei that have different roles in affect regulation ^25^, we further assessed the independent role of the centromedial and laterobasal amygdala groups. Second, we used effective connectivity as a measure of directed or causal connectivity between areas ^26^ to assess the influence of the amygdala, insula and vmPFC on one another. We also included the NAc in this model because 1) DBS was targeted at the border of the NAc and vALIC, 2) the NAc is strongly connected with the amygdala, insula, and vmPFC ^21^, and 3) we previously observed DBS related changes in NAc connectivity ^11^. We used spectral dynamic causal modelling (DCM)^27,28^, which is particularly suited to measure group differences in effective connectivity during the resting state. It is based on constructing a biologically plausible model which generates a predicted response in the frequency domain and fitting that to the observed response. This enables one to infer causal influences one region exerts over another. DCM provides estimation of parameters that give information on the strength of those causal influences between regions of interest (ROIs), referred to as effective connectivity ^27,29^. To test the influence of DBS on these parameters, patients were investigated twice. The first resting state scan was obtained after DBS treatment for at least a year (DBS on), and the second resting state scan was obtained when stimulation was turned off for one week (DBS off). To control for test‐retest effects on the connectivity measures, a group of healthy controls was also measured twice. Based on the positive association between anxiety and amygdala‐insula connectivity and the negative association between amygdala‐vmPFC connectivity, we hypothesized that effect of DBS treatment on mood and anxiety could either be explained by decreased amygdala‐insula connectivity, increased amygdala‐vmPFC connectivity, or both.

## Methods

### Participants

Sixteen patients with treatment refractory OCD were recruited from the outpatient clinic for deep brain stimulation at the Department of Psychiatry of the Academic Medical Center in Amsterdam (AMC), the Netherlands. Symptom severity was assessed using the Yale‐Brown Obsessive‐Compulsive Scale (Y‐BOCS) ^30,31^, the Hamilton Depression Rating Scale (HAM‐D)^32^ and the Hamilton Anxiety Rating Scale (HAM‐A) ^33^. Two quadripolar electrodes (Model 3389 (Medtronics Inc., Minneapolis, USA) with 4 contact points of 1.5 mm long and intersected by 0.5 mm spaces were implanted bilaterally through the ALIC with the deepest contact point located in the nucleus accumbens (NAc) in the plane 3 mm anterior to the anterior commisure and the 3 upper contact points positioned in the vALIC. Patients were included only if they had undergone an optimization phase of at least one year where they were evaluated every 2 weeks for severity of symptoms during which the stimulation parameters were adjusted accordingly. For all patients the optimal stimulation was monopolar using the two middle contact points superior to the NAc, in the ventral ALIC. We could not collect all resting state fMRI data for 3 patients, 1 patient had a deviating electrode placement, and data from 2 patients were excluded due to excessive head motion during scanning (max movement > 2.5 mm or 2.5 degree of rotation), leaving a final sample for data analysis of 10 patients. At the commencement of the study the mean stimulation voltage was 4.8 Volt (3.5‐6.2V), frequency was 130 Hz (9 patients) or 185 Hz (1 patients). The pulse width was 90 microseconds (7 patients) or 150 microseconds (3 patients). We recruited 16 healthy control participants from the community via local advertisements. Exclusion criteria were the presence of a mental disorder according to DSM‐IV as assessed with the Mini International Neuropsychiatric Inventory (MINI) ^34,35^, a family history of psychiatric disease, a history of head trauma, any neurological or other medical disorders, a history of substance abuse, or a contraindication for MRI. Data from one session of one patient was missing and data of four controls were excluded due to excessive head motion during scanning, leaving a final sample size of 11 controls. During the two scanning days, the participants did not use cigarettes, caffeine or sedatives. The study was approved by the Medical Ethics Committee of the AMC and all participants signed an informed consent form before participation.

### Study design

Each subject underwent two resting state fMRI scans. The first scan (DBS ON) was performed after DBS had been constantly turned on for at least 1 year. After the first scan, patients entered the DBS off phase, and the second scan was performed one week later (DBS OFF). Symptom severity was assessed on each scanning day through the use of Y‐BOCS, HAM‐D and HAM‐A. The healthy controls were scanned with one week in between sessions.

### Image acquisition

Data was acquired using a 1.5T Siemens MAGNETOM Avanto scanner. A transmit receive head coil was used to minimize exposure of DBS electrodes to the pulsed radiofrequency field. The DBS was turned off 2 min prior to scanning and programmed at 0V in bipolar mode. Head of subject was held in place with padding and straps. Specific absorption rate (SAR) was limited to 0.1 W kg^‐1^. Structural images were acquired with 1×1×1 mm resolution using a 3D sagittal MPRAGE with repetition time (TR) of 1.9 s, echo time (TE) of 3.08 ms, flip angle of 8° and inversion time (TI) of 1.1 s. FMRI data were acquired with two dimensional echo‐planar imaging with TR = 2000 ms, TE = 30 ms, field angle = 90°. Each scan consisted of 25 transverse slices of 4 mm thick with in plane voxel size of 3.6×3.6 mm and slice gap of 0.4 mm. The first ten volumes were discarded to allow for magnetization stabilization and the subsequent 180 volumes were analyzed.

### Image Preprocessing

Image processing was carried out using Nipype, a pipeline tool for neuroimaging data processing ^36^ and SPM12 (http://www.fil.ion.ucl.ac.uk/spm). The functional data were first realigned to correct for motion using rigid body realignment to the first functional image. Subjects exhibiting more than 2.5 mm movement in any direction were excluded, resulting in 10 patients and 11 controls. After motion correction the brain extracted structural data was coregistered to functional space. The structural data was normalized to MNI152 space using SPM’s unified segmentation approach ^37^ that is robust to brain lesions ^38^ to accommodate the signal dropout due to the DBS system. The resulting nonlinear warps were then used to normalize the functional data, which were resampled into 2 mm isotropic voxels and subsequently smoothed using an 8mm Gaussian kernel and bandpass filtered from 0.01 to 0.1 Hz.

### Functional connectivity analysis

Data from regions of interest (ROI) were extracted using SPM’s Anatomy toolbox ^39^. For each amygdala, the centromedial (CM) and laterobasal (LB) sub regions were extracted. The mean signal in the ROI was computed from the unsmoothed but bandpass filtered functional data. For the sub regions the mean signal was weighted with the probability at each voxel as was done previously ^40^. This gives stronger weight to those voxels more probable to belong to each sub region. Noise correction followed the procedure in ^41^. Principal components were extracted from the cerebrospinal fluid (CSF) and white matter (WM) signals. The CSF tissue mask from SPM’s unified segmentation was confined to the ventricles using the ALVIN mask ^42^ and including only voxels with a 99% probability or higher as being CSF. The WM mask was confined to a 99% probability or higher and eroded to minimize the risk of capturing signals from the nearby gray matter regions. For the signal at each voxel the voxel mean was removed and the results divided by the voxel standard deviation. Then singular value decomposition (SVD) was used to generate principal components. The components accounting for the 50% of variance from each tissue class were included in the nuisance regression as covariates along with the 6 motion parameters and their derivatives (computed using backward differences). The motion parameters were bandpass filtered to prevent inadvertent reintroduction of nuisance related variation into frequencies previously suppressed by bandpass filtering ^43^. For each analysis, the other sub region from the same hemisphere was included in the nuisance regression. After nuisance covariates were regressed out the ROI signal was correlated with the remaining residuals from the whole brain. This provided a map of correlation coefficients which were then transformed to Z scores using Fisher’s transformation.

The individual statistical maps were entered into a 2×2 flexible factorial design using GLM Flex (http://mrtools.mgh.harvard.edu/) with partitioned error terms for the within and between subject factors. The factors included were Group (patients vs controls) and Condition (DBS ON vs OFF). All main effects and interactions were computed. Voxel‐wise statistical tests were family wise error (FWE) corrected for multiple comparisons at the cluster level (p<0.05) using a cluster forming threshold of p=0.001 ^44^ using peak_nii (https://www.nitrc.org/projects/peak_nii) for the whole brain or the regions of interest (ROI). The insula was extracted from the automatic anatomical labelling atlas ^45^ using wfu Pickatlas ^46^ and the vmPFC was defined as a 10 mm sphere centered on (‐1, 49, ‐5) MNI coordinates ^47^. Post‐hoc t‐tests were performed in presence of significant interactions between factors. Data were extracted from the entire ROI for brain regions that showed significant clusters for subsequent correlation analyses with clinical scores.

### Dynamic causal modelling

Because functional connectivity analyses do not provide information about the direction of connectivity we subsequently performed an effective connectivity analysis using spectral DCM. First a general linear model was set up in SPM with cosine basis functions from 1/128 Hz to 0.1 Hz as effects of interest and the movement parameters, white and grey matter signal as nuisance regressors. This way the resulting effects of interest contrast over the basis functions reveals the resting state fluctuations in that frequency range. For model simplicity only ROIs from the left hemisphere were included since the functional connectivity results were with the left amygdala. We included the amygdala and insula as ROIs but also the NAc and vmPFC, because all these regions are anatomically connected ^21^ and we previously showed that vALIC DBS also influences NAc vmPFC connectivity ^11^.The left amygdala was specified based on an anatomical atlas using the SPM anatomy toolbox. The NAc was defined as the caudate nucleus from the automatic anatomical atlas ^39^ below z=0 and excluding the signal dropout due to the DBS electrodes. The vmPFC ROI was specified as a 10mm sphere centered on (‐1, 49, ‐5) MNI coordinates ^47^. The left insula ROI was specified as a 10 mm sphere centered on the peak value in the ROI from the seed‐based correlation analysis. When extracting the signal, the spheres were allowed to shift to the nearest local maxima according to the effects of interest contrast defined above, but within the 10 mm sphere. From each ROI the first principal eigenvariate, corrected for confounds, was used to represent the ROI.

A DCM model was constructed with the four ROIs as nodes. Bilateral connections between all nodes were defined, resulting in 16 connections including each node’s self‐connection (see Figure 1).

**Figure 1:**
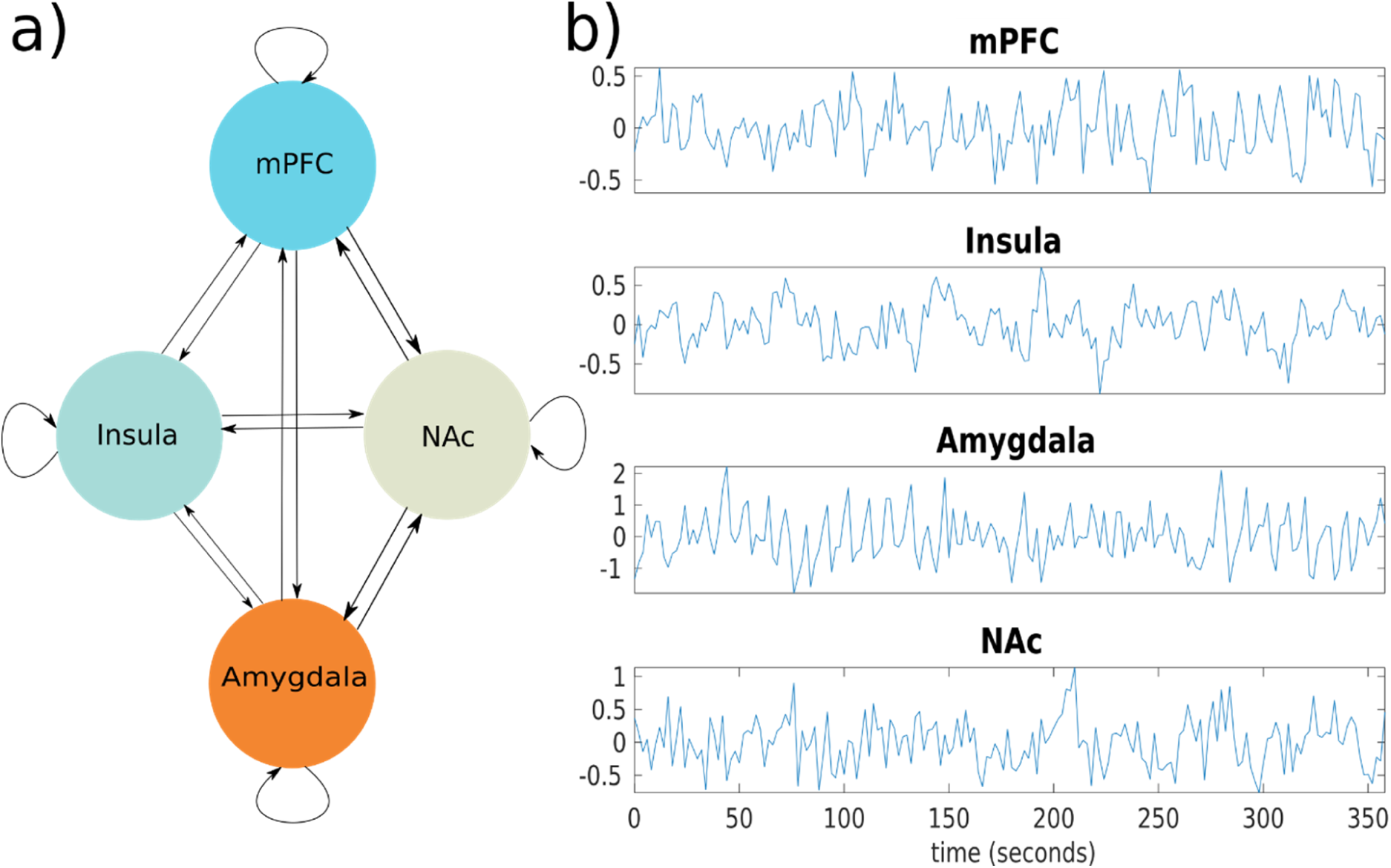
The causal neural model for the effects of DBS. A) A graph model showing the fully connected model with four regions, the mPFC, insula, amygdala, and Nac. In a fully connected model each region is reciprocally connected and each region has a self‐inhibitory connection. B) Bold time series data from regions of interest from one subject.

Then for each subject this full model was inverted using spectral DCM in SPM (DCM12 revision 6662). The resulting posterior probabilities of each subject’s connection coefficients were then entered into a repeated measures mixed ANOVA in Matlab and solved for significant interactions between DBS ON and OFF in patients and controls.

## Results

### DBS effects on mood and anxiety

Demographics and clinical characteristics are summarized in Table 1. All participants were right‐handed. The patients and controls did not differ in age, sex ratio, years of education and head motion during fMRI scanning (all p>0.05; table 1). All patients had OCD as primary diagnosis, four patients had comorbid major depressive disorder, one had comorbid panic disorder, and three had comorbid obsessive‐compulsive personality disorder. In line with our previous report on the clinical outcome of DBS ^9^, turning off DBS increased anxiety symptoms (HAM‐A; t(9) = 2.84, p=0.019, paired t‐test), increased mood symptoms (HAM‐D; t(9)=3.31,p=0.009, paired t‐test), and increased obsessive‐compulsive symptoms (Y‐BOCS; t(9) = 3.46, p = 0.007).

**Table 1:** demographics of the study sample and clinical scales

|  | Patients (n=10) |  | Controls (n=11) |  | Difference |
| --- | --- | --- | --- | --- | --- |
|  | Mean (SD) | Range | Mean (SD) | Range | P-value |
| Age, y. | 44.1 (9.7) | 27-56 | 44.7 (9.1) | 25-56 | 0.88<sup>1</sup> |
| Gender (% Males). | 50 |  | 63.6 |  | 0.37<sup>2</sup> |
| Education, y. | 13.7 (1.56) | 12-16 | 15.7 (4.07) | 12-23 | 0.16<sup>1</sup> |
| Smoking (% Yes). | 30 |  | 54.5 |  | 0.76<sup>2</sup> |
| Illness Duration, y. | 27.6 (13.4) | 8-48 |  |  |  |
| Motion (mean framewise displacement), mm. | 0.26 (0.13) | 0.09-0.55 | 0.21 (0.08) | 0.09-0.36 | 0.16 |

| Clinical scales patients | DBS OFF |  | DBS ON |  | P-value |
| --- | --- | --- | --- | --- | --- |
|  | Mean (SD) | Range | Mean (SD) | Range |  |
| Y-BOCS total | 28.5 (6.3) | 15-38 | 18.9 (7.7) | 6-32 | $< 0.01^{3*}$ |
| Y-BOCS obsessions | 13.7 (3.3) | 9-19 | 9.0 (4.1) | 0-15 | $< 0.01^{3*}$ |
| Y-BOCS compulsions | 14.8 (4.0) | 6-20 | 9.9 (4.0) | 6-18 | $< 0.05^{3*}$ |
| HAM-D | 30 (9.2) | 13-40 | 16.7 (10.4) | 0-30 | $< 0.01^{3*}$ |
| HAM-A | 38.3 (11.3) | 11-51 | 17.7 (9.2) | 4-31 | $< 0.05^{3*}$ |

Abbreviations: Y-BOCS: Yale-Brown Obsessive-Compulsive Scale; HAM-D: Hamilton Rating Scale for Depression; HAM-A: Hamilton Rating Scale for Anxiety
<sup>1</sup>independent sample t-test. <sup>2</sup> Chi-square test. <sup>3</sup>Paired t-test
\*Significant (2 tailed)

### The impact of DBS on functional connectivity of the amygdala

Functional connectivity analysis showed a significant interaction between group (patients vs controls) and session (DBS ON vs OFF) of the left LB amygdala groups with the right insula (p=0.014, mni: (44, ‐2, 4), size: 248mm^3^, FWE corrected, figure 1). Left amygdala functional connectivity with the right insula increased from DBS ON to DBS OFF in OCD patients, whereas it decreased between session in controls. Post‐hoc tests showed that the interaction was primarily driven by an increase in LB amygdala‐insula connectivity when DBS was switched OFF in the patient groups (p=0.009). Further, connectivity tended to be higher in patients than controls when DBS was switched OFF (p=0.078), and was not significantly different when DBS was ON (p=0.51).

The DBS‐induced change in connectivity was positively correlated to changes in anxiety (r=0.67, p=0.035) and mood (r=0.67, p=0.033), such that a larger DBS‐related increase in LB amygdala‐insula connectivity was associated with higher increase in mood and anxiety symptoms. The correlation between DBS‐induced changes in mood and anxiety symptoms was 0.9 (p=0.0004), precluding further partial correlations and suggesting that the influence of DBS on these symptoms cannot be dissociated.

### Dynamic causal modeling for DBS effects

Our analysis showed an interaction between group and session for the connection between the left vmPFC and left amygdala (p=0.045, FDR corrected), and between the left amygdala and left insula (p=0.045, FDR corrected). The impact of the vmPFC over the amygdala was higher during DBS ON compared to DBS OFF in patients (p=0.0122) and was higher during DBS ON compared to controls (p=0.0012). The connection parameter for the impact of amygdala on insula was positive during DBS OFF, indicating an excitatory connection, whereas it was negative during DBS ON, indicating an inhibitory connection (p=0.0052). Moreover, the connection parameter was lower in patients during DBS ON compared to controls (p=0.033, figure 2).

**Figure 2.**
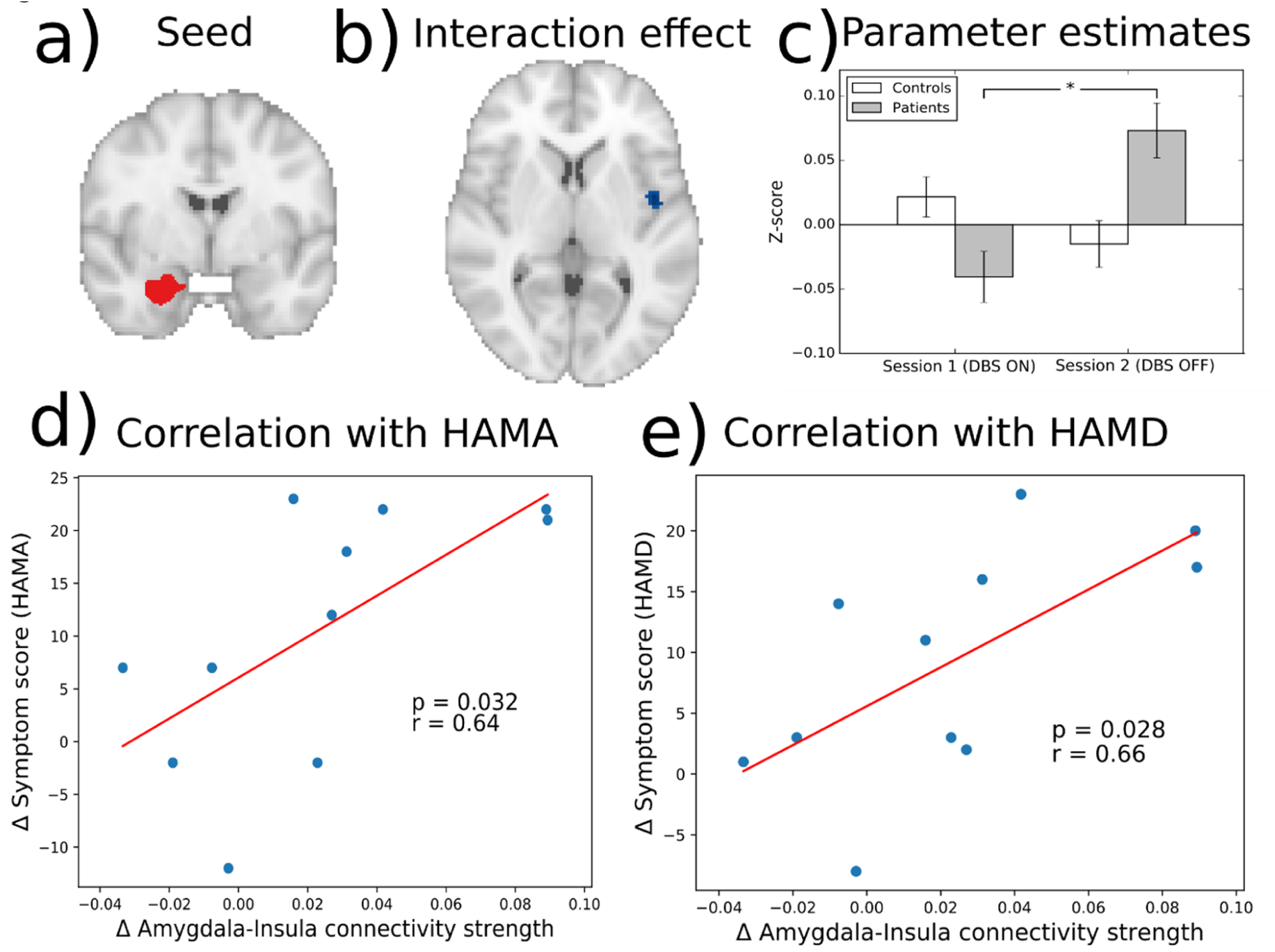
The effects of DBS on functional connectivity of the laterobasal amygdala with the insula. a) The left LB amygdala seed region. b) The significant interaction cluster in the right insula. c) Parameter estimates (±standard error) for the significant interaction cluster (for illustrative purposes). d) Correlation between changes in laterobasal amygdala‐insula connectivity and HAM‐A scores in OCD patients. The blue dots are data from each patient while the red line is a fitted regression line. E) Correlation between changes in laterobasal amygdala‐insula connectivity and HAM‐D scores in OCD patients. *p<0.05

**Figure 3.**
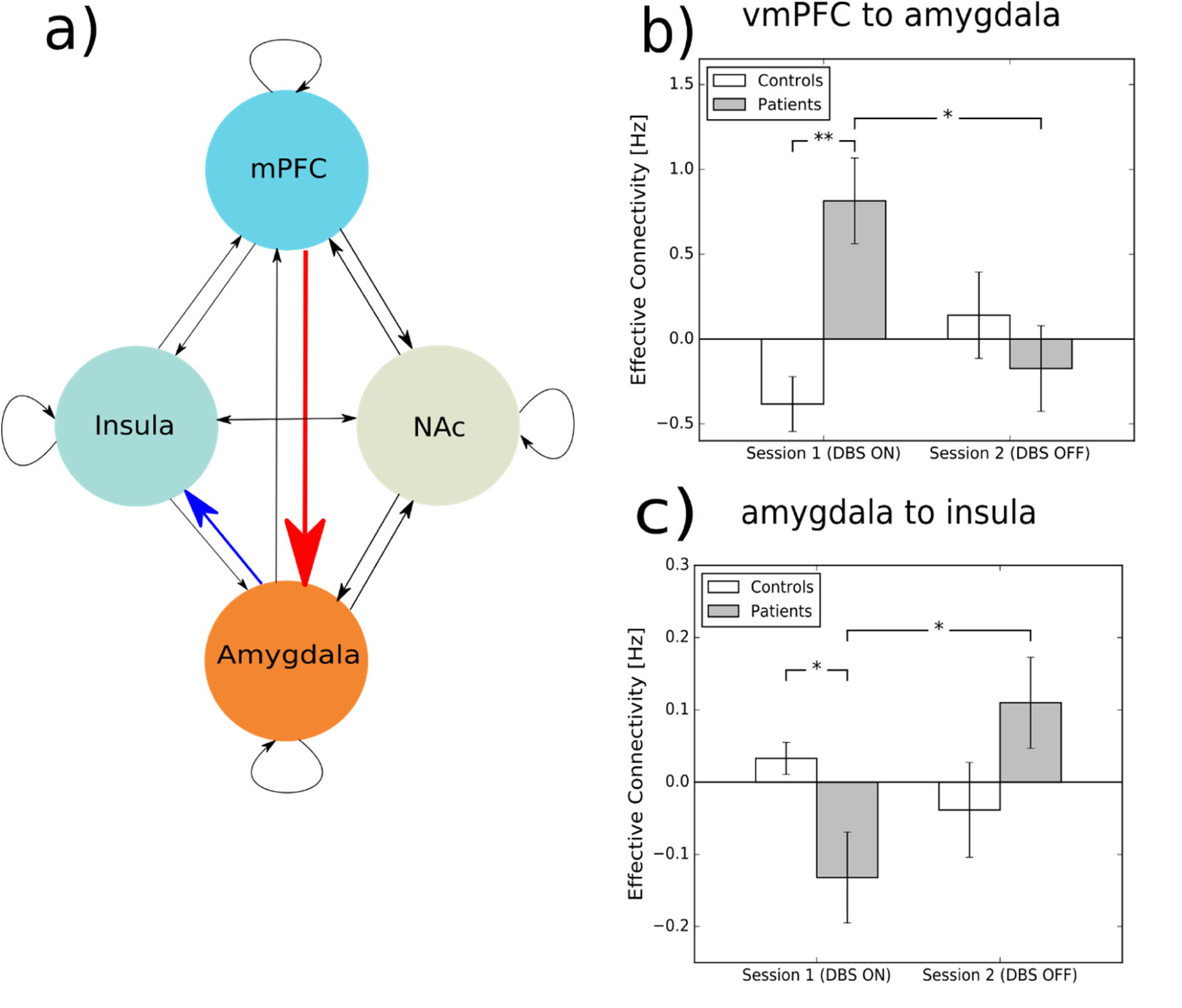
The influence of DBS on effective connectivity between the amygdala, insula, vmPFC, and Nac. A) A graph model showing the fully connected model with four regions, the mPFC, insula, amygdala, and Nac. In a fully connected model, each region is reciprocally connected and each region has a self‐inhibitory connection. b) The connection from vmPFC to the amygdala is higher when the DBS is on and is higher than in controls. c) The connection from amygdala to insula reverses direction from inhibitory influence to excitatory when DBS is turned off. Error bars are standard errors. *p<0.05, **p<0.001

## Discussion

When undergoing vALIC DBS for OCD, the most prominent and rapid changes in symptoms are improvements in mood and anxiety. To better understand the underlying mechanism, we investigated functional connectivity within the amygdala network and found that turning DBS off increased functional connectivity between amygdala and insula. This increase was correlated to an increase of both anxiety and mood symptoms. When analyzing directional connectivity within the network, we found that during DBS the impact of vmPFC on amygdala was higher than when DBS was switched OFF and turning DBS OFF reversed the impact of amygdala on insula from inhibitory to excitatory.

This affective prefrontal‐limbic network has primarily been linked to mood and anxiety disorders ^13,14,17–19^, but it is also associated with OCD. In particular, OCD‐symptom provoking stimuli induce exaggerated amygdala responses ^16,48,49^. It has been suggested that elevated fear and anxiety, and associated fronto‐limbic impairments, may be causal to, or driving some of the compulsions ^50^. This is supported by a recent meta‐analysis ^51^ that found increased activation of the amygdala during emotional processing in OCD, and that comorbidity with mood and anxiety disorders was associated with even higher activations of the right amygdala, putamen and insula as well as lower activations in the left amygdala and right vmPFC. The insula is involved in perception of internal feelings ^52^ and is suggested to have an important role in anxiety ^53^. In OCD, the insula has been primarily associated with disgust sensitivity ^54^, but animal studies suggest that it also has a crucial role in the development of compulsive behavior (Belin‐Rauscent et al., 2015). Our results suggest that the effects of vALIC DBS on mood and anxiety are primarily driven by changes in connectivity between insula and amygdala, demonstrating the importance of these brain regions in OCD symptomatology.

Further, we showed that DBS alters the top‐down control of vmPFC on amygdala and reverses amygdala drive on insula from excitatory to inhibitory. It has been shown that the strength of amygdala coupling with the vmPFC predicts the extent of attenuation of negative affect by reappraisal in healthy subjects ^55^, indicating that normal top‐down control of the amygdala can lower anxiety and negative mood. OCD patients show reduced vmPFC‐amygdala coupling during appraisal and passive viewing of symptom stimuli ^56^. In addition, reduced activity in the vmPFC is also associated with impaired recall of fear extinction in OCD ^57^. The retention of extinction memory is crucial for the success of extinction training, and is therefore also thought to underlie the success of exposure therapy. So the change in top‐down control of the vmPFC on the amygdala might mean that DBS restores reappraisal mechanisms and facilitates fear extinction, and can thereby improve mood and anxiety to enable successful cognitive behavioral therapy ^10^.

The current pathophysiological model for OCD is centered around hyperactivity in cortico‐striatal‐thalamic loops and does not fully explain negative mood and anxiety. In line with previous suggestions ^50^, our results suggest that abnormal frontolimbic connectivity needs to be incorporated into that model. In fact the temporal sequence of symptom changes following DBS shows that improvements in mood and anxiety happen before improvements in obsessions and compulsions ^9^. The changes in frontolimbic connectivity might therefore precede and enable the changes in frontostriatal circuits that are associated with obsessive‐compulsive symptoms ^11^. The influence of DBS on the frontolimbic circuit may also underlie the restoration of self‐confidence, as is typically reported by patients. We recently hypothesized that in particular anxiety fuels low self‐confidence in OCD, which could be related to insufficient vmPFC control over the amygdala ^58^.

Frontolimbic network has primarily been implicated in other anxiety disorders and depression ^13^. vALIC DBS is also beneficial for treatment resistant depression, suggesting that DBS‐induced changes in vmPFC‐amygdala‐insula connectivity might also be important for its clinical effects in major depression ^59^. At the same this, this implies that vALIC DBS might be beneficial for other treatment‐resistant anxiety disorders for which DBS is not yet a treatment option. Besides DBS, similar neural network changes may underlie clinical improvement induced by other treatment options such as pharmacotherapy and psychotherapy, which could be explored in future studies.

There are a few limitations to our study. First the sample size is small as can be expected for a neuroimaging study with fully implanted DBS electrodes, which previously has only been investigated in several cases ^60^. The small sample size did not allow us to assess the influence of clinical heterogeneity or concurrent medication use. Second, spectral DCM is a technique which is hard to validate in absence of known causal network changes, however it has been shown to be able to recover those changes in synthetic data ^28^ and was found to be more sensitive to group differences than conventional DCM. Further, a recent study shows that spectral DCM has good inter‐subject and inter‐session reliability when studying the default mode network ^61^. Third, a relatively small number of nodes can be used in spectral DCM due to computational reasons, requiring a priori specification of regions of interest. We therefore only selected those nodes which had shown DBS‐related changes in functional connectivity in the current and a previous study ^11^. Fourth, in our study we cannot disentangle the effects of DBS on mood or anxiety separately due to the high correlation between them. Further work is needed to see if DBS is affecting one more than the other or if mood and anxiety change concurrently.

In conclusion, our results reveal a neural network model for how vALIC DBS could exert its rapid effects on mood and anxiety, which may enable patients to challenge their obsessive‐compulsive symptoms with behavioral therapy. Mood and anxiety symptoms may therefore be more important in OCD than often appreciated, or at least they are critical symptoms targeted by effective vALIC DBS for OCD. In fact, the initial modulation of the frontolimbic circuit may enable later alterations in the frontostriatal circuit, which we found is related to eventual changes in compulsivity ^11^. Beyond OCD, the frontolimbic network also has an important role in other anxiety disorders and depression ^13,17^. This suggests that modulation of the vmPFC‐amygdala‐insula circuit may also have a role in the clinical effects of vALIC DBS in depression ^59^ and highlights its potential as novel treatment option for patients with other treatment‐resistant anxiety disorders. Future studies may investigate whether other treatment modalities exert their antidepressant and anxiolytic effects through similar neural mechanisms.

## Acknowledgments

This study was supported by an unrestricted investigator‐initiated research grant by Medtronic Inc (Denys and Schuurman), which provided the devices used herein. The sponsor had no role in the design or conduct of the study; in the collection, management, analysis, or interpretation of the data; or in the preparation, review, or approval of the manuscript..

## Author contributions

M.F., D.D. and G.v.W. designed the study. E.A.F. performed all the analyses. P.v.d.M. and P.R.S. performed neurosurgeries. M.F. and J.L. conducted functional neuroimaging. E.A.F., G.v.W. and D.D. prepared the manuscript. All authors contributed to discussions about the results and critically revised the manuscript.

## References

1. Godlewska BR, Norbury R, Selvaraj S, Cowen PJ, Harmer CJ. Short-term SSRI treatment normalises amygdala hyperactivity in depressed patients. Psychol Med. 2012;42(12):2609–2617. doi:10.1017/S0033291712000591

2. Ruscio AM, Stein DJ, Chiu WT, Kessler RC. The epidemiology of obsessive-compulsive disorder in the National Comorbidity Survey Replication. Mol Psychiatry. 2010. doi:10.1038/mp.2008.94

3. Denys D. Pharmacotherapy of Obsessive-compulsive Disorder and Obsessive-Compulsive Spectrum Disorders. Psychiatr Clin North Am. 2006;29(2):553–584. doi:10.1016/j.psc.2006.02.013

4. Alonso P, Cuadras D, Gabriels L, et al. Deep Brain Stimulation for Obsessive-Compulsive Disorder: A Meta-Analysis of Treatment Outcome and Predictors of Response. PLoS One. 2015;10(7):e0133591. doi:10.1371/journal.pone.0133591

5. J. Luis Lujan, Ashutosh Chaturvedi CCM. Tracking the mechanisms of deep brain stimulation for neuropsychiatric disorders. Front Biosci. 2008. doi:10.2741/3124

6. Luigjes J, de Kwaasteniet BP, de Koning PP, et al. Surgery for Psychiatric Disorders. World Neurosurg. 2013;80(3-4):S31.e17–S31.e28. doi:10.1016/J.WNEU.2012.03.009

7. de Koning PP, Figee M, van den Munckhof P, Schuurman PR, Denys D. Current Status of Deep Brain Stimulation for Obsessive-Compulsive Disorder: A Clinical Review of Different Targets. Curr Psychiatry Rep. 2011;13(4):274–282. doi:10.1007/s11920-011-0200-8

8. Coenen VA, Schlaepfer TE, Goll P, et al. The medial forebrain bundle as a target for deep brain stimulation for obsessive-compulsive disorder. CNS Spectr. 2017;22(3):282–289. doi:10.1017/S1092852916000286

9. Denys D, Mantione M, Figee M, et al. Deep brain stimulation of the nucleus accumbens for treatment-refractory obsessive-compulsive disorder. Arch Gen Psychiatry. 2010;67(10):1061–1068.

10. Mantione M, Nieman DH, Figee M, Denys D. Cognitive–behavioural therapy augments the effects of deep brain stimulation in obsessive–compulsive disorder. Psychol Med. 2014;44(16):3515–3522. doi:10.1017/S0033291714000956

11. Figee M, Luigjes J, Smolders R, et al. Deep brain stimulation restores frontostriatal network activity in obsessive-compulsive disorder. Nat Neurosci. 2013;16(4):386–387. doi:10.1038/nn.3344

12. Davis M, Whalen PJ. The amygdala: vigilance and emotion. Mol Psychiatry. 2001;6(1):13–34. doi:10.1038/sj.mp.4000812

13. Taylor JM, Whalen PJ. Neuroimaging and Anxiety: the Neural Substrates of Pathological and Non-pathological Anxiety. Curr Psychiatry Rep. 2015;17(6). doi:10.1007/s11920-015-0586-9

14. Etkin A, Wager TD. Functional neuroimaging of anxiety: A meta-ana lysis of emotional processing in PTSD, social anxiety disorder, and specific phobia. Am J Psychiatry. 2007;164(10):1476–1488. doi:10.1176/appi.ajp.2007.07030504

15. Via E, Cardoner N, Pujol J, et al. Amygdala activation and symptom dimensions in obsessive-compulsive disorder. Br J Psychiatry. 2014;204(1):61–68. doi:10.1192/bjp.bp.112.123364

16. Simon D, Adler N, Kaufmann C, Kathmann N. Amygdala hyperactivation during symptom provocation in obsessive-compulsive disorder and its modulation by distraction. NeuroImage Clin. 2014;4:549–557. doi:10.1016/j.nicl.2014.03.011

17. Hamilton JP, Etkin A, Furman DJ, Lemus MG, Johnson RF, Gotlib IH. Functional neuroimaging of major depressive disorder: a meta-analysis and new integration of base line activation and neural response data. Am J Psychiatry. 2012;169(7):693–703. doi:10.1176/appi.ajp.2012.11071105

18. Baur V, Hänggi J, Langer N, Jäncke L. Resting-State Functional and Structural Connectivity Within an Insula–Amygdala Route Specifically Index State and Trait Anxiety. Biol Psychiatry. 2013;73(1):85–92. doi:10.1016/j.biopsych.2012.06.003

19. Kim MJ, Gee DG, Loucks RA, Davis FC, Whalen PJ. Anxiety Dissociates dorsal and ventral medial prefrontal cortex functional connectivity with the amygdala at rest. Cereb Cortex. 2011;21(7):1667–1673. doi:10.1093/cercor/bhq237

20. Morawetz C, Bode S, Baudewig J, Heekeren HR. Effective amygdala-prefrontal connectivity predicts individual differences in successful emotion regulation. Soc Cogn Affect Neurosci. 2017;12(4):569–585. doi:10.1093/scan/nsw169

21. Cho YT, Ernst M, Fudge JL. Cortico-Amygdala-Striatal Circuits Are Organized as Hierarchical Subsystems through the Primate Amygdala. J Neurosci. 2013;33(35):14017–14030. doi:10.1523/JNEUROSCI.0170-13.2013

22. Zuo XN, Kelly C, Adelstein JS, Klein DF, Castellanos FX, Milham MP. Reliable intrinsic connectivity networks: Test-retest evaluation using ICA and dual regression approach. Neuroimage. 2010;49(3):2163–2177. doi:10.1016/j.neuroimage.2009.10.080

23. Shehzad Z, Kelly AMC, Reiss PT, et al. The resting brain: Unconstrained yet reliable. Cereb Cortex. 2009;19(10):2209–2229. doi:10.1093/cercor/bhn256

24. Clare Kelly AM, Uddin LQ, Biswal BB, Castellanos FX, Milham MP. Competition between functional brain networks mediates behavioral variability. Neuroimage. 2008;39(1):527–537. doi:10.1016/j.neuroimage.2007.08.008

25. Herry C, Johansen JP. Encoding of fear learning and memory in distributed neuronal circuits. Nat Neurosci. 2014;17(12):1644–1654. doi:10.1038/nn.3869

26. Friston KJ. Functional and effective connectivity: a review. Brain Connect. 2011;1(1):13–36. doi:10.1089/brain.2011.0008

27. Friston KJ, Kahan J, Biswal B, Razi A. A DCM for resting state fMRI. Neuroimage. 2014;94:396–407. doi:10.1016/j.neuroimage.2013.12.009

28. Razi A, Kahan J, Rees G, Friston KJ. Construct validation of a DCM for resting state fMRI. Neuroimage. 2015;106. doi:10.1016/j.neuroimage.2014.11.027

29. Friston KJ, Harrison L, Penny W. Dynamic causal modelling. Neuroimage. 2003;19(4):1273–1302. doi:10.1016/S1053-8119(03)00202-7

30. Goodman WK, Price LH, Rasmussen S a, et al. The Yale-Brown Obsessive Compulsive Scale. I. Development, use, and reliability. Arch Gen Psychiatry. 1989;46(11):1006–1011. doi:10.1001/archpsyc.1989.01810110048007

31. Goodman WK, Price LH, Rasmussen SA, et al. The Yale-Brown Obsessive Compulsive Scale. II. Validity. Arch Gen Psychiatry. 1989;46(11):1012–1016. doi:10.1001/archpsyc.1989.01810110048007

32. Hamilton M. A Rating Scale for Depression. J Neurol Neurosurg Psychiat. 1960;23:56–62. doi:10.1136/jnnp.23.1.56

33. Hamilton M. Hamilton Anxiety Rating Scale (HAM-A). J Med. 1959;61(4):81–82. doi:10.1145/363332.363339

34. Sheehan D V., Lecrubier Y, Sheehan KH, et al. The Mini-International Neuropsychiatric Interview (M.I.N.I.): The development and validation of a structured diagnostic psychiatric interview for DSM-IV and ICD-10. In. Journal of Clinical Psychiatry. Vol 59.; 1998:22–33. doi:10.1016/S0924-9338(99)80239-9

35. van Vliet IM, de Beurs E. [The MINI-International Neuropsychiatric Interview. A brief structured diagnostic psychiatric interview for DSM-IV en ICD-10 psychiatric disorders]. Tijdschr Psychiatr. 2007;49(6):393–397. doi:TVPart_1639 [pii]

36. Gorgolewski K, Burns CD, Madison C, et al. Nipype: a flexible, lightweight and extensible neuroimaging data processing framework in python. Front Neuroinform. 2011;5(August):13. doi:10.3389/fninf.2011.00013

37. Ashburner J, Friston KJ. Unified segmentation. Neuroimage. 2005;26(3):839–851. doi:10.1016/j.neuroimage.2005.02.018

38. Crinion J, Ashburner J, Leff A, Brett M, Price C, Friston K. Spatial normalization of lesioned brains: Performance evaluation and impact on fMRI analyses. Neuroimage. 2007;37(3):866–875. doi:10.1016/j.neuroimage.2007.04.065

39. Eickhoff SB, Stephan KE, Mohlberg H, et al. A new SPM toolbox for combining probabilistic cytoarchitectonic maps and functional imaging data. Neuroimage. 2005;25(4):1325–1335. doi:10.1016/j.neuroimage.2004.12.034

40. Roy AK, Shehzad Z, Margulies DS, et al. Functional connectivity of the human amygdala using resting state fMRI. Neuroimage. 2009;45(2):614–626. doi:10.1016/j.neuroimage.2008.11.030

41. Muschelli J, Nebel MB, Caffo BS, Barber AD, Pekar JJ, Mostofsky SH. Reduction of motion-related artifacts in resting state fMRI using aCompCor. Neuroimage. 2014;96:22–35. doi:10.1016/j.neuroimage.2014.03.028

42. Kempton MJ, Underwood TSA, Brunton S, et al. A comprehensive testing protocol for MRI neuroanatomical segmentation techniques: Evaluation of a novel lateral ventricle segmentation method. Neuroimage. 2011;58(4):1051–1059. doi:10.1016/j.neuroimage.2011.06.080

43. Hallquist MN, Hwang K, Luna B. The nuisance of nuisance regression: Spectral misspecification in a common approach to resting-state fMRI preprocessing reintroduces noise and obscures functional connectivity. Neuroimage. 2013;82:208–225. doi:10.1016/j.neuroimage.2013.05.116

44. Eklund A, Nichols TE, Knutsson H. Cluster failure: Why fMRI inferences for spatial extent have inflated false-positive rates. Proc Natl Acad Sci. 2016. doi:10.1073/pnas.1602413113

45. Tzourio-Mazoyer N, Landeau B, Papathanassiou D, et al. Automated Anatomical Labeling of Activations in SPM Using a Macroscopic Anatomical Parcellation of the MNI MRI Single-Subject Brain. Neuroimage. 2002;15(1):273–289. doi:10.1006/nimg.2001.0978

46. Maldjian JA, Laurienti PJ, Kraft RA, Burdettea JH. An automated method for neuroanatomic and cytoarchitectonic\ratlas-based interrogation of fMRI data sets. Neuroimage. 2003;19(3):1233–1239. doi: Doi 10.1016/S1053-8119(03)00169-1

47. Fox MD, Snyder AZ, Vincent JL, Corbetta M, Van Essen DC, Raichle ME. The human brain is intrinsically organized into dynamic, anticorrelated functional networks. Proc Natl Acad Sci U S A. 2005;102(27):9673–9678. doi:10.1073/pnas.0504136102

48. Simon D, Kaufmann C, Müsch K, Kischkel E, Kathmann N. Fronto-striato-limbic hyperactivation in obsessive-compulsive disorder during individually tailored symptom provocation. Psychophysiology. 2010. doi:10.1111/j.1469-8986.2010.00980.x

49. Van Den Heuvel OA, Veltman DJ, Groenewegen HJ, et al. Amygdala activity in obsessive-compulsive disorder with contamination fear: A study with oxygen-15 water positron emission tomography. Psychiatry Res - Neuroimaging. 2004. doi:10.1016/j.pscychresns.2004.06.007

50. Milad MR, Rauch SL. Obsessive-compulsive disorder: Beyond segregated cortico-striatal pathways. Trends Cogn Sci. 2012. doi:10.1016/j.tics.2011.11.003

51. Thorsen AL, Hagland P, Radua J, et al. Emotional Processing in Obsessive-Compulsive Disorder: A Systematic Review and Meta-analysis of 25 Functional Neuroimaging Studies. Biol Psychiatry Cogn Neurosci Neuroimaging. 2018. doi:10.1016/j.bpsc.2018.01.009

52. Craig AD. How do you feel - now? The anterior insula and human awareness. Nat Rev Neurosci. 2009. doi:10.1038/nrn2555

53. Paulus MP, Stein MB. An Insular View of Anxiety. Biol Psychiatry. 2006. doi:10.1016/j.biopsych.2006.03.042

54. Shapira NA, Liu Y, He AG, et al. Brain activation by disgust-inducing pictures in obsessive-compulsive disorder. Biol Psychiatry. 2003. doi:10.1016/S0006-3223(03)00003-9

55. Banks SJ, Eddy KT, Angstadt M, Nathan PJ, Luan Phan K. Amygdala-frontal connectivity during emotion regulation. Soc Cogn Affect Neurosci. 2007. doi:10.1093/scan/nsm029

56. Paul S, Beucke JC, Kaufmann C, et al. Amygdala-prefrontal connectivity during appraisal of symptom-related stimuli in obsessive-compulsive disorder. Psychol Med. 2019;49(2):278–286. doi:10.1017/S003329171800079X

57. Milad MR, Furtak SC, Greenberg JL, et al. Deficits in conditioned fear extinction in obsessive-compulsive disorder and neurobiological changes in the fear circuit. JAMA Psychiatry. 2013. doi:10.1016/B978-0-08-097086-8.25066-6

58. Kiverstein J, Rietveld E, Slagter HA, Denys D. Obsessive Compulsive Disorder: A Pathology of Self-Confidence. Trends Cogn Sci. 2019. doi:10.1016/j.tics.2019.02.005

59. Bergfeld IO, Mantione M, Hoogendoorn MLC, et al. Deep Brain Stimulation of the Ventral Anterior Limb of the Internal Capsule for Treatment-Resistant Depression. JAMA Psychiatry. 2016;73(5):456. doi:10.1001/jamapsychiatry.2016.0152

60. Rauch SL, Dougherty DD, Malone D, et al. A functional neuroimaging investigation of deep brain stimulation in patients with obsessive–compulsive disorder. J Neurosurg. 2006. doi:10.3171/jns.2006.104.4.558

61. Almgren H, Van de Steen F, Kühn S, Razi A, Friston K, Marinazzo D. Variability and reliability of effective connectivity within the core default mode network: A multi-site longitudinal spectral DCM study. Neuroimage. 2018;183:757–768. doi:10.1016/j.neuroimage.2018.08.053

